# Fungal growth is affected by and affects pH and redox potential (Eh) of the growth medium

**DOI:** 10.1101/401182

**Authors:** L. Bousset, M. Ermel, B. Soglonou, O. Husson

## Abstract

Fungal plant pathogens live in the specific environment of plants. Understanding of the interaction between pathogens and their host plants might open new ways to control plant diseases. Yet, the specificities of the plant environment and its effects on fungal growth are not yet fully explored. Both pH and Eh play a key role during the interaction between the fungus and its host plants, but often studied independently or at different scales. To decipher the interrelation between plant growth and their soil environment, recent theoretical and methodological advances have been made through the joint characterization of the pH and Eh. This opens the prospect to develop similar methods for fungi. The aim of our study was to investigate whether the methods developed for soil could be transposed to fungi. We first worked on artificial media, assessing the impact of fungal growth on the media in cultures. The growth of all 16 species tested significantly altered either Eh, pH or both in agar media. Measuring Eh reveals that even the species not modifying pH can have an impact on the surrounding environment. Reciprocally, altered media were used to characterize sensitivity of fungal growth to both pH and Eh parameters. The response of the six fungi tested to the modified media was quantitative with a decrease in colony diameter. In addition, colony aspect was repeatedly and thoroughly modified. As a first step towards the same studies in conditions matching the natural environment of fungal pathogens, we tested how the measurement can be performed with fungi growing on oilseed rape plants. In infected plant stems, pH and Eh were significantly altered, in opposite directions for *L. maculans* and *S. sclerotiorum*. The observed alcalinisation or acidification correlates with canker length. Our series of experiments indicate that the procedure published for Eh and pH in soil can be extended for measurement in agar media and in infected plants. Further, the joint characterization of both parameters opens the way to a more precise understanding of the impact of fungi on their environment, and conversely, of the environment on fungal growth. The availability of methods for measurement opens the prospect to study combinations of stresses, either on agar media or in plants, and get an understanding of the involvement of pH and Eh modifications in these interactions.

## Introduction

Fungal plant pathogens need to get resources from the plant and for doing so they need to live in the specific environment of plants, with an host species ranging from broad for generalists to narrow for specialist (Braunsdorf *et al*., 2016; Zeilinger *et al*., 2016; van der Does & Rep, 2017). Getting a better understanding of the interaction between pathogens and their host plants might open new ways to control plant diseases. Within species, the interaction between fungal pathogens and plant resistance has been approached through molecular dialog between partners (Lo Presti *et al*., 2015; Jwa & Hwang, 2017). Yet, the specificities of the plant environment and its effects on fungal growth are not yet fully explored. To decipher the interrelation between plant growth and their soil environment, recent theoretical and methodological advances have been made through the joint characterization of the pH and Eh (Husson *et al*., 2013; 2018). This opens the prospect to develop similar methods for fungi, because both pH and Eh are important in their metabolism.

As compared to other microorganisms, fungi are capable of growth and development over wide pH ranges. Several species, including plant pathogens, can actively modulate the pH of their environment by secreting acids or alkali (Landraud *et al*., 2013; Vylkova *et al*., 2017). The ability to control extracellular pH is an important aspect of fungal physiology that contributes to fitness within the host (Vylkova *et al*., 2017) via the modulation of virulence factors to best fit the host (Prusky & Yakoby, 2003). pH is revealed as one component at work during interaction between the fungus and its host plants, from field to molecular scales. At the field scale, opposite effect can be observed for wheat soilborne diseases: take-all (*Gaeumannomyces graminis var. tritici*) is more severe in alkaline soils (Smiley, 1974; Lebreton *et al*., 2014) and cephalosporium stripe (*Cephalosporium gramineum*) severity increase with soil acidity (Stiles & Murray, 1996). Soil pH is considered as a major structuring ambient factor of telluric microbial communities and plant microbiome (Rousk *et al*., 2010; Chapelle *et al*., 2016). In culture, pH is taken into account for the optimization of biocoltrol agents production as *Trichoderma* species (Singh *et al*., 2014). In plant tissues, pH indicators or image analysis have been proposed to monitor changes and fungal growth (Tardi-Ovadia *et al*., 2017). At the molecular level, pH alters gene expression (Daval *et al*., 2013). All these elements indicate that for fungi, pH is an important characteristic of the environment. It has however been stressed that because of their interrelation, analyses of pH gain to be associated with measures of Eh, that is oxido-reduction status (Husson et al. 2016; 2018).

So far, in plant-pathogenic fungi interactions, oxido-reduction processes are documented mostly at cellular and molecular levels (Lehman *et al*., 2015; Jwa & Hwang, 2017). Plants have been investigated more thoroughly. Oxido-reduction reactions and reactive oxygen species have a profound influence on almost every aspect of plant biology especially to balance information from metabolism and the environment (Dietz *et al*., 2016; Foyer & Noctor, 2016). The concept that reactive oxygen species (ROS) and reactive nitrogen species (RNS) are key signaling molecules that facilitate a plethora of adaptive metabolic, molecular genetic and epigenetic responses is now established (Foyer & Noctor, 2005; 2016). The balance between oxidant and antioxidant pool sizes, plays signaling roles in the regulation of gene expression and protein function in a wide variety of plant physiological processes including stress acclimation (Shigeoka & Maruta, 2014; Choudhury *et al*., 2017), regulation of plant development processes, organogenesis, senescence and defense against pathogens (Nosek *et al*., 2015; Marschall & Tudzynski, 2016; Noctor *et al*., 2017). As oxido-reduction processes play a major role in plant biology, developing methods to investigate if this occurs also in fungi is desirable.

Eh and pH are interacting, as oxidation leads to acidification, and the importance of measuring both parameters has been documented on soils (Husson *et al*., 2018). Eh and pH regulation is central in plant physiology and phenology and as the vast majority of biotic and abiotic stresses are translated into redox signals in plants (Husson, 2013). The leaf Eh and pH, were proposed in previous studies to explain the variable resistance of cereals to obligates parasites (Benada, 2017). Specifically, these studies investigated the disease gradients on plants; the change of susceptibility of organs during the ontogeny and growth; the difference in resistance in individual plant cells and the rapid change of resistance in a couple of hour (Benada, 2017). Investigating these topics further could benefit from the recent methodological developments. The aim of our study was to investigate whether the methods developed for soil (Husson *et al*., 2016; 2018) could be transposed to fungi. We first worked on artificial media, assessing the impact of fungal growth on the media in fungal cultures. Reciprocally, altered media were used to characterize sensitivity of fungal growth to both pH and Eh parameters. Later, we tested how the measurement can be performed with fungi growing on oilseed rape plants, as a first step towards the same studies in conditions matching the natural environment of fungal pathogens.

## Materials and Methods

### Culture media

To test how fungal growth modified the medium on which they grew, four independent experiments were performed with 6 fungal strains (Table 1). In the Mar2016 experiment, 13 additional strains were included (Table 1). All tests were performed on malt-agar media (Malt extract 20 g.l^-1^, Agar 20 g.l^-1^, Streptomycin 0.1 g.l^-1^). All culture media were autoclaved for 20mn at 120°C and cooled down before Streptomycin concentrated solution 10% w/v in water was added. Within each experiment, replicates were prepared independently and corresponding uninoculated aliquots were kept for pH and Eh initial and final measurements (Table 1). Fungal cultures were grown in sterile 6 cm diameter plastic containers (50 ml) commonly used to store and reheat food sauce.

**Table 1:**
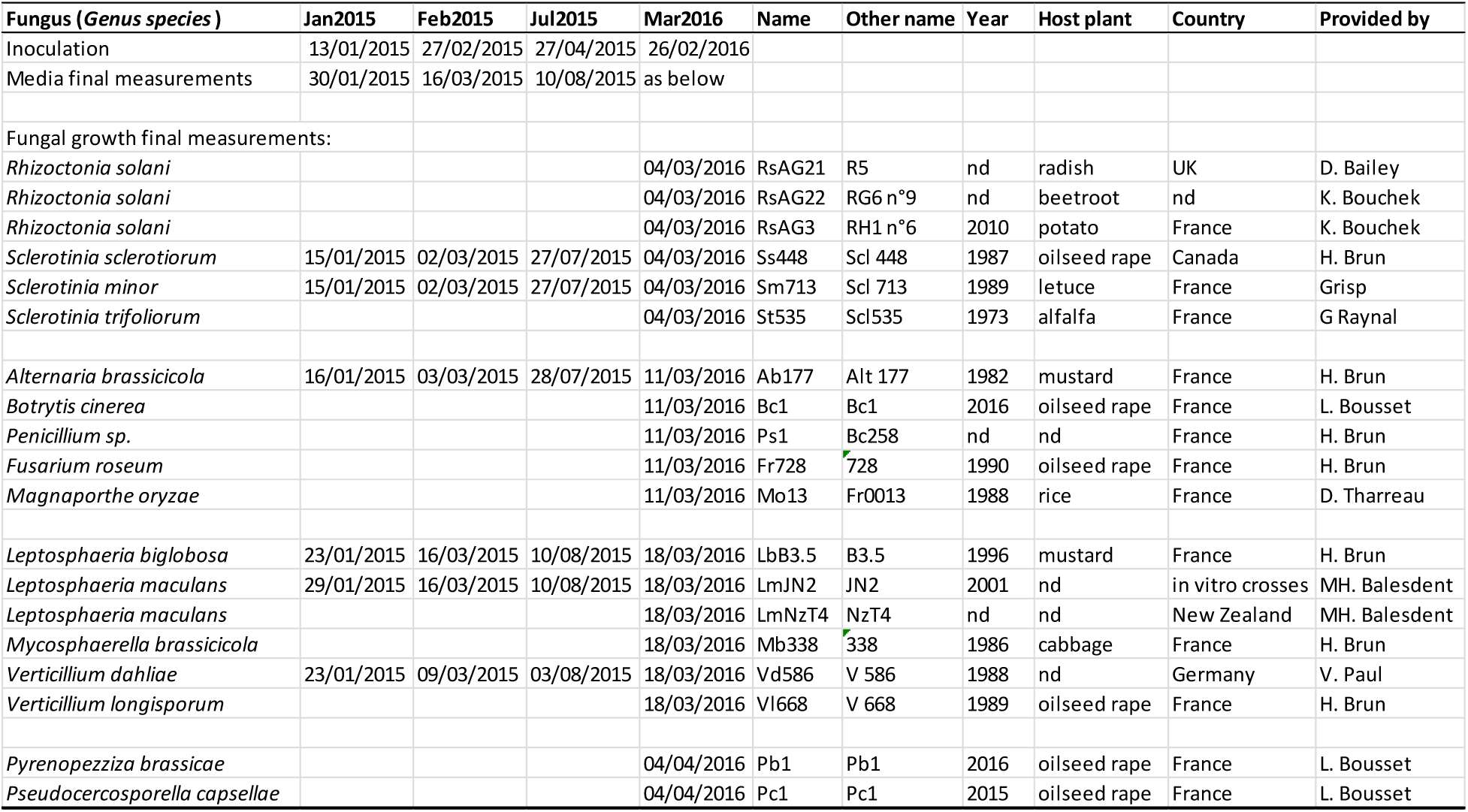
List and origin of 19 fungal strains used in the 4 experiments (Jan2015 to Mar2016) with dates of inoculation, final measurements for media and fungal growth. “nd” stands for unknown.

To test how modified pH and Eh media altered fungal growth, within each of the 3 experiments (Jan2015, Feb2015, Jul2015) three replicates of a range of 13 malt-agar media (Malt extract 20 g.l^-1^, Agar 20 g.l^-1^, Streptomycin 0.1 g.l^-1^) were prepared independently. To alter Eh and pH without changing the composition of the media, we used electrolyzed water products, because when chemical compounds are added, it is difficult to disentangle the effects via pH from directs effects (Bekker *et al*., 2009). The range of agar media included native one (N0) and four sets of three media with part of the water (in proportions 1:3; 1:1 and 3:1) replaced by either of catholyte (pH 11;7, 11.5, 12.0 in the three experiments), basic anolyte (pH 8.1, 8.7, 9.0), neutral anolyte (pH 7.1, 7.1, 7.3) and acid anolyte (pH 3.3, 2.0, 1.5). Aliquots were prepared in 20 ml flacons for uninoculated measurements at the start and end of the experiment. Aliquots of 10 ml in 5 cm diameter Petri dishes were prepared for inoculation and fungal growth measurement. For each experiment, anolyte and catholyte were obtained from IDEEAQUACULTURE, Parc Euromédecine 2, 39 Rue Jean Giroux, 34080 Montpellier, France. Products were freshly prepared 24 to 48h before media preparation, using Envirolyte ELA900 and shipped in light-opaque containers. Eh and pH of the products were measured on aliquots before use. These products are all obtained from water, and the only compound added during the production is NaCl. We thus made a specific experiment (Sep2015, see below) to check that within the range of final NaCl concentrations in the media, fungal growth was not altered for any of the 6 fungal species (Fig. S1).

To test how modified pH and Eh media altered fungal growth of *M. oryzae*, within two experiments (Jan2016, Mar2016) four replicates of a range of 13 rice flour-agar media (Rice flour 10 g.l^-1^, Agar 15 g.l^-1^, Yeast extract 2 g.l^-1^, Penicillin 0.1 g.l^-1^) were prepared. The range of rice flour agar media included native one (N0), addition of Hydrogen peroxide, addition of Hydroquinone and the combination of both. This made 20 media in Jan2016 with combinations (Hydroquinone - Hydrogen peroxide) 0-0; 0- 0.05; 0-0.1; 0-0.2; 0-0.4; 0-0.8; 0-1.6; 4-0; 4-0.1; 16-0; 16-0.05; 16-0.1; 16-0.2; 16-0.4; 64-0; 64-0.1; 256-0; 256-0.05; 256-0.1; 256-0.4 and 14 media in Mar2016 with combinations (Hydroquinone – Hydrogen peroxide) 0-0; 0-1; 0-2; 0-4; 50-0; 50-2; 100-0; 100-1; 100-2; 100-4; 20-0; 200-2; 300-0; 300-4.

### Fungal growth experiments

A total of 19 isolates of 16 fungal species were used, either from the INRA Le Rheu collection or kindly provided by colleagues (Table 1). For each isolate, 5 mm diameter agar plugs of fresh cultures on malt-agar were prepared. Agar plug were placed in the center of Petri dishes and fungal growth was measured as two perpendicular diameters of the colony (Table 1). To assess the impact of fungal growth on the medium, plastic containers were used instead of Petri dishes in order to allow sufficient depth for the electrodes after colony growth. Agar plug was placed in the center, and final measurements were performed by sticking electrodes through the colony, on both sides of the plug. In each of the 3 replicates, there was 1 Petri dish and 1 container in Jan2015 and Feb2015; 2 Petri dishes in Sep2015; 3 Petri dishes and 3 containers in Jul2015 and 3 containers in Mar2016 experiments, respectively. All cultures and uninoculated aliquots were kept at room temperature in INRA Le Rheu, with no supplementary light. Hourly temperature (mean 21.7°C se 1.6) and humidity (mean 46.5% se 8.9) were recorded with a TEMPCO Hygro bouton 23 data logger.

*M.oryzae* growth experiments were performed at Africarice, Cotonou, Bénin with local strains BN1162 in Jan16 and BN1168 in Mar16. For each isolate, 5 mm diameter agar plugs of fresh cultures on rice flour-agar were prepared. Agar plug were placed in the center of Petri dishes and fungal growth was measured as two perpendicular diameters of the colony.

### Oxido-reduction Potential (ORP) electrodes and voltmeter

Redox potential was measured in agar media as Described by Husson *et al*. (2016) using Ag/AgCl Reference electrode Radiometer analytical E21M002 and Radiometer Analytical Pt plate electrode 5×5 mm M241 Pt, with a Voltcraft VC850 multimeter (10×10^6^ Ohm input resistance) for all fungi and a WTW 3110 (10×10^12^ Ohm input resistance) for *M. oryzae* growth experiments. The Eh value considered was the Eh (in mV) indicated on the meter after 1 minute without change in the Eh unit. After being measured according to Ag/AgCl reference electrode, all redox potentials are transformed to give Eh according to the Normal Hydrogen Electrode (ENH). Redox electrodes were calibrated at the start of the measurements and every 10-12 measurements, with Mettler Toledo Redox buffer solution 220 mV (pH 7) composed of Potassium hexacyanoferrate (III), Potassium hexacyanoferrate (II), Potassium dihydrogen phosphate and Disodium hydrogen phosphate. All measurements were conducted outdoor, in an environment identified as being free of electromagnetic interference.

pH was measured in agar media with a with a Voltcraft VC850 multimeter (10×10^6^ Ohm input resistance) using Ag/AgCl reference electrode Radiometer analytical E21M002 and a glass pH electrode Radiometer analytical pHG301-9. pH electrodes were calibrated at the start of the measurements and every 10-12 measurements, with VWR buffer solutions at pH4, pH7 and pH10. Data were converted into pH values using linear regression of the measured values of standard solutions.

### Plants experiments

Winter oilseed rape was sown in 03/10/2017 in 12 x 12 x 21 cm pots with a 1:1:1 mix of sand, peat and compost and grown outside at INRA Le Rheu (48·1°N, 1·5°W), in Brittany, France. The experiment matches the usual cultivation in this area, where winter oilseed rape is generally sown in late August – September and harvested the following July. There was three replicates of 38 plants for *L. maculans* experiments and 2 replicates of 38 and 44 plants for *S. sclerotiorum* experiments.

Natural infections with *L. maculans* airborne ascospores occurred on some of the plants, with typical leaf spots observed in November. Cankers became visible on some stems at flowering. Measurements were performed on infected and uninfected plants after flowering, over 4 days (02/05; 07/05; 22/05, 28/05). Plant stem was cut lengthwise, and internal canker length was recorded. Immediately after, a 2 cm portion of the stem taken at crown level was cut and quickly grinded with a mortar and pestle. The plant material was then placed in the barrel of a 2 ml syringe from which the nozzle had been cut. For the redox potential measurement with the Voltcraft VC850 multimeter, we used a 9 cm Petri dish with a filter paper moistened with 0.1M KCl solution (Fig. S2). The Ag/AgCl reference electrode was placed standing, with tip touching the moist filter paper. The Pt plate electrode was placed in the plant material in the syringe, then the syringe barrel was placed standing with tip touching the moist filter paper. After reading the redox potential, the plunger of the syringe was placed in the barrel to squeeze the liquid onto a Horiba LAQUAtwin-pH-22 meter for pH measurement.

Inoculations with *S. sclerotiorum* were performed at the time of petal fall. Matchsticks pieces were autoclaved in liquid 2% malt medium. Agar plugs from fresh culture were transferred (13/04 and 30/04) to the center of 9 cm diameter Petri dishes (Malt extract 20 g.l-1, Agar 20 g.l-1, Streptomycin 0.1 g.l-1) with lying matchsticks pieces, and cultured at 20°C. On 17/04 and 03/05 for each of the 2 plant replicates, a matchstick piece covered with mycelium was tightly fitted through the 3 mm diameter hole drilled at 10 cm height through the stem of 16 and 24 inoculated and 19 and 23 mock-inoculated plants. External canker length was measured 20/04 and 07/05, immediately after, a 2 cm portion of the stem taken at the inoculation point was cut and processed for Eh and pH measurement as described above.

## Data analysis

Statistical analyses on artificial media experiment data were performed using XLSTAT (Addinsoft, 2016). Analysis of variance (ANOVA) was conducted on Eh, pH for comparison of agar media after fungal growth. ANOVAs were followed by a comparison of means using the Ryan-Einot-Gabriel-Welsh test Q (REGW-Q) as it provides the best compromise between the need to have a powerful test and the need to limit the familywise error rate at α, according to Howell (2009). Quadratic regressions between colony growth, pH and Eh were performed for the different 7 fungal species on altered agar media.

Statistical analyses on plant experiment data were performed using R software (R Core Team, 2013). Generalized linear modelling was used to investigate the effects of inoculation, canker on pH measured. For plant pH and Eh data, we used a Gaussian linear model on log-transformed data. Linear contrasts were calculated with as Estimated Marginal Means with emmeans package to determine specific differences among treatment combinations.

## Results

### Effect of fungi on pH and Eh in the agar media

Uninoculated malt-agar media were acid (pH from 4.5 to 5) and oxidized (Eh from 500 to 600) (Fig. 1). For all tested species, ANOVA analyses followed by REGW-Q comparison of means indicated that fungal growth significantly altered pH and Eh in agar media (Fig. 1; Table 2). These alterations were species-specific and in directions opposite to the slight evolution of uninoculated media between initial (empty symbols) and final (filled symbols) measurements. All *Sclerotinia* species acidified the media, *S. minor* and *S. sclerotiorum* did not change Eh, while *S. trifolorium* also reduced the media (Fig. 1A). *M. oryzae, Penicillium sp*. and *P. brassicae* alcalinised the media, with variable intensities of reduction. The main effect of other species was reduction, either mild (Bc1, Mb, Pc) intermediate (Lm, Vd) or intense (Rs, Ab, Lb, Fr). Results of independent experiments with 6 species were congruent (Fig. 1B). The Eh-pH couples obtained after the growth of the fungal species tested did not align on the theoretical correlation between Eh and pH, with a negative slope of 59mV decrease in Eh for one unit increase in pH at 25°C, according to the equation of Nernst (Fig. 1AB).

**Table 2:**
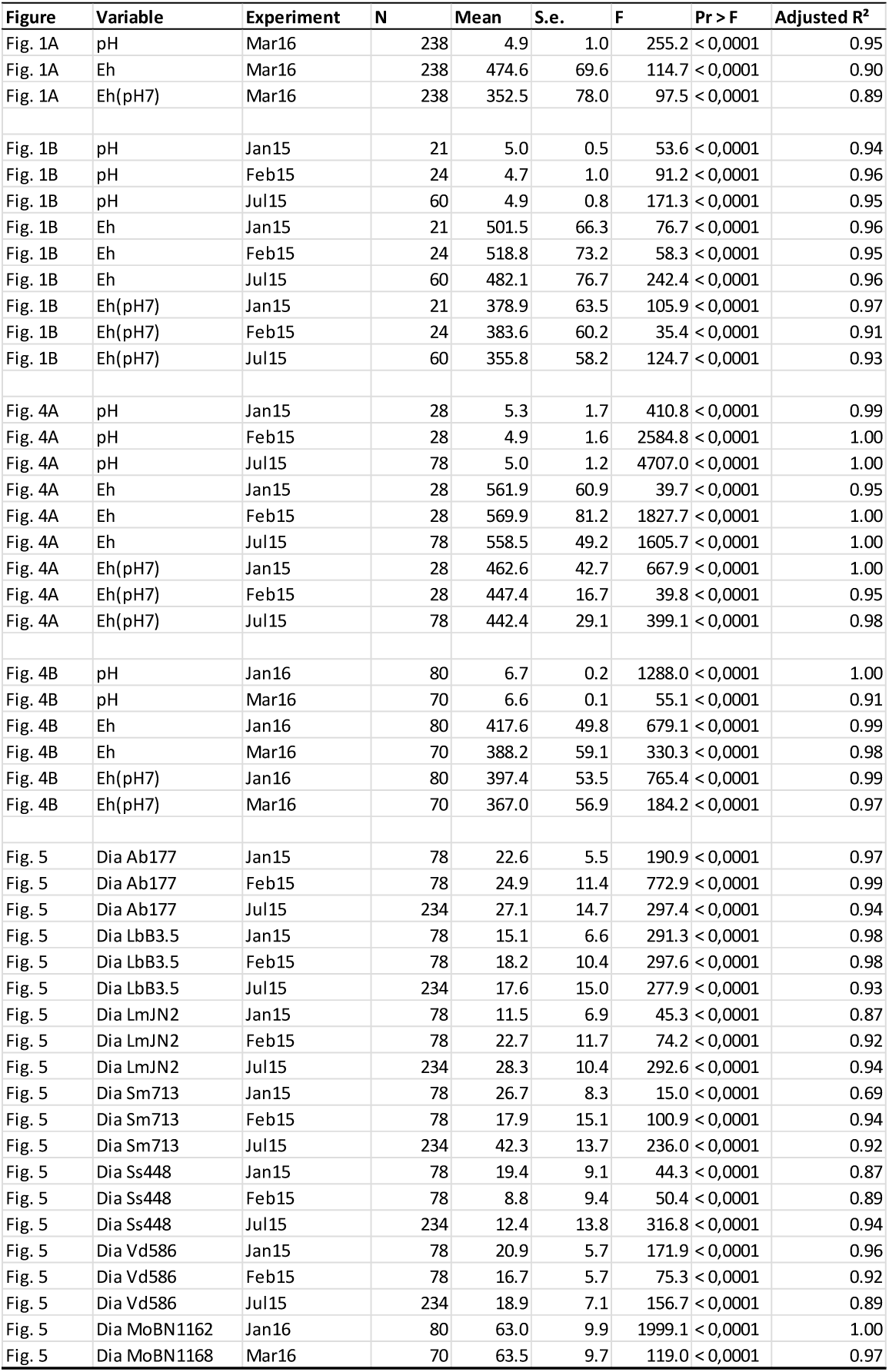
For each figure and each experiment, mean and standard error of the variable are indicated, with F and p-values of the ANOVA. Significantly different groups in the REGW-Q comparisons are added as dashed lines on the figures.

**Fig. 1.**
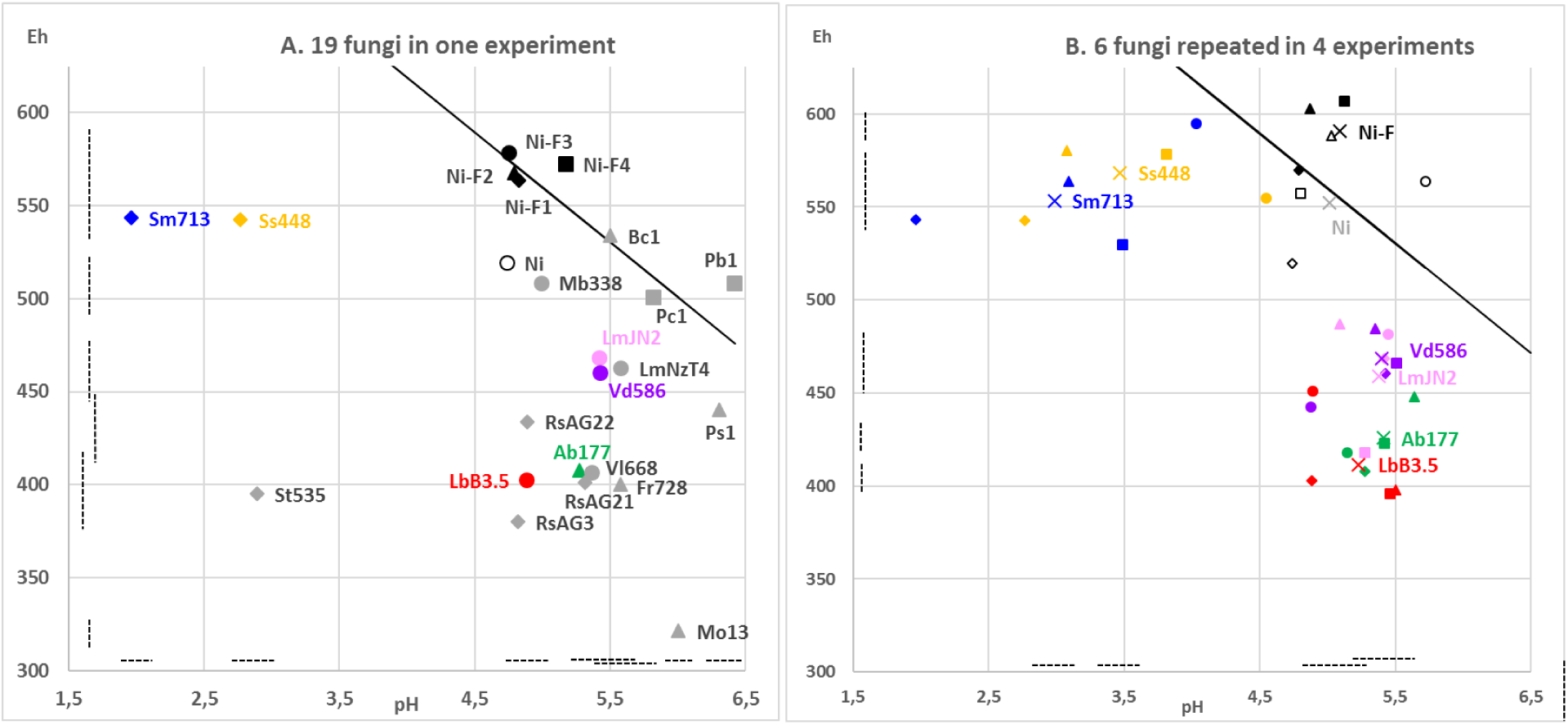
Effect of fungal growth on the pH and Eh of the malt-agar media A. for 19 fungi in March16 experiment; B. for 6 fungi in 4 independent experiments of Jan2015 (diamonds) Feb2015 (triangles) Jul2015 (squares) and Mar2016 (circles) with overall means (crosses). Fungi in panel B are *S. minor* (blue) *S. sclerotiorum* (yellow) *L. maculans* (pink) *L. biglobosa* (red) *A. brassicicola* (green) *V. dahliae* (purple) and the remaining ones (grey) in panel A are detailed in Table 1. Uninoculated medium (black) was measured at the start (Ni, empty symbols) and at the end (Ni-F, filled symbols) of the experiments. Symbols in Panel A correspond to measurement at different final dates (see Table 1). Each point is mean of 2 measures in each of 3 containers in Jan15 and Fev15, 9 containers in Jul15 and Mar16. Plain line shows the theoretical relationship between pH and Eh according to the equation of Nernst. Dashed lines correspond to significantly different groups for pH (horizontal) and Eh (vertical) in the REGW-Q analysis after ANOVA (see Table 2).

### Effect of fungi on pH and Eh in plant stems

Uninfected oilseed rape stems were slightly acidic (pH from 5.5 to 6.5) like malt-agar media, but more reduced (Eh from 150 to 300) as compared to malt-agar media (Eh from 500 to 600).

Analysis of deviance showed that both inoculation status and canker length were significant predictors of changes in pH (Table 3), though in opposite directions depending on the fungal species. pH was higher following fungal infection with *L. maculans* (Fig. 2A; Table 3) whereas pH was lower with *S. sclerotiorum* (Fig. 3A; Table 3). For both fungi, canker length had a significant effect on measured pH (Fig. 2B; Fig. 3B, Table 3).

**Fig. 2.**
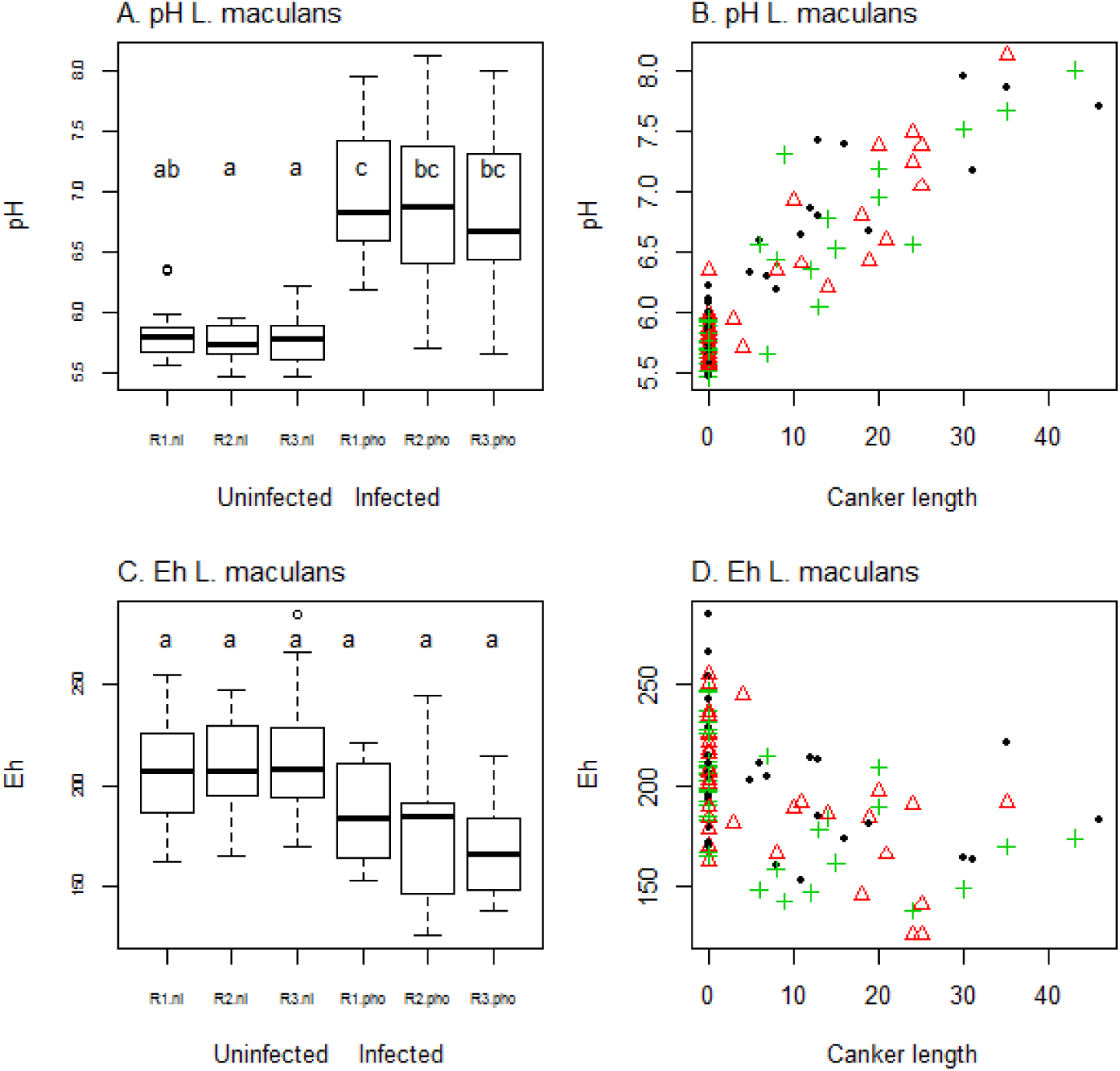
Effect of fungal growth on the pH and Eh of oilseed rape stems. A. Boxplot of pH at stem base on plants uninfected or infected with *L. maculans* for the three replicates (R1, R2, R3). B. Measured pH in stems depending on canker length, symbols differ for the three sets of plants. C. Boxplot of Eh at stem base on plants uninfected or infected with *L. maculans* for the three replicates (R1, R2, R3). B. Measured Eh in stems depending on canker length, symbols differ for the three sets of plants.

**Fig. 3.**
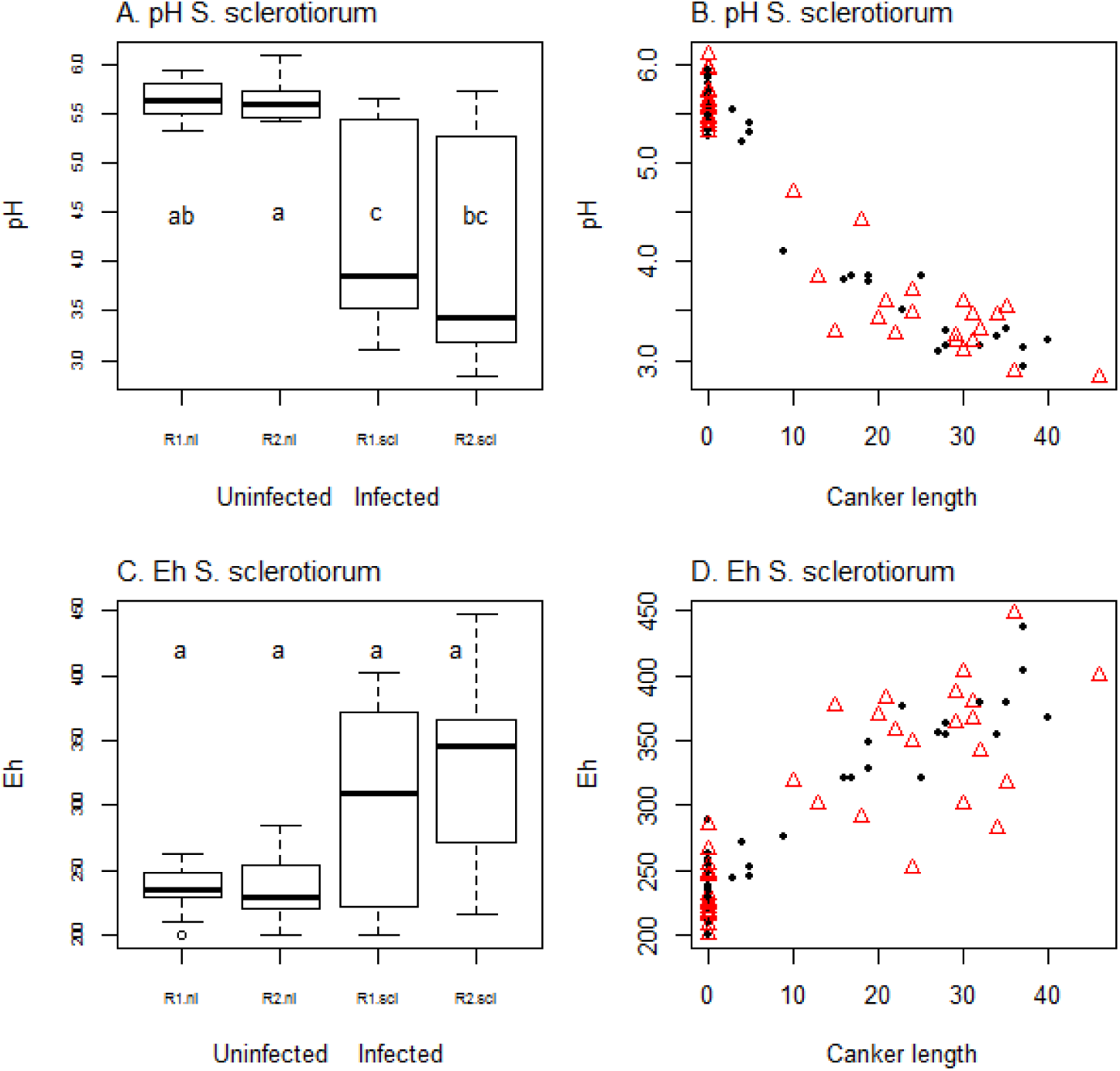
Effect of fungal growth on the pH and Eh of oilseed rape stems. A. Boxplot of pH at stem base on plants uninfected or infected with *S. sclerotiorum* for the two replicates (R1, R2). B. Measured pH in stems depending on canker length, symbols differ for the two replicates. C. Boxplot of Eh at stem base on plants uninfected or infected with *S. sclerotiorum* for the two replicates (R1, R2, R3). B. Measured Eh in stems depending on canker length, symbols differ for the two replicates.

**Table 3.** Analysis of deviance on logN transformed data (Type II tests). Pr(χ2) are p-values from a χ2 test used for significance

| Model | Tested effect | Df | $\chi^2$ | Pr(>$\chi^2$) | |
| --- | --- | --- | --- | --- | --- |
| *L. maculans* pH | Canker | 1 | $1.28 \times 10^2$ | $< 2 \times 10^{-16}$ | *** |
| | Infection | 2 | $1.48 \times 10^5$ | $< 2 \times 10^{-16}$ | *** |
|  | Replicate | 2 | 2.41 | 0.30 |  |
| *L. maculans* Eh | Canker | 1 | 2.51 | 0.11 |  |
| | Infection | 2 | $1.26 \times 10^5$ | $< 2 \times 10^{-16}$ | *** |
|  | Replicate | 2 | 0.56 | 0.75 |  |
| *S. sclerotiorum* pH | Canker | 1 | 616.64 | $< 2 \times 10^{-16}$ | *** |
| | Infection | 2 | 32608.66 | $< 2 \times 10^{-16}$ | *** |
|  | Replicate | 1 | 3.01 | 0.08 | . |
| *S. sclerotiorum* Eh | Canker | 1 | $1.75 \times 10^2$ | $< 2 \times 10^{-16}$ | *** |
| | Infection | 2 | $1.30 \times 10^5$ | $< 2 \times 10^{-16}$ | *** |
|  | Replicate | 1 | 2.65 | 0.10 |  |

Analysis of deviance showed that inoculation status was a significant predictor of changes in Eh (Table 3), again in opposite directions depending on the fungal species. Eh was lower following fungal infection with *L. maculans* though differences between groups were not significant (Fig. 2C; Table 3) whereas Eh was higher with *S. sclerotiorum* (Fig. 3C; Table 3). The effect of canker length on Eh was significant for *S. sclerotiorum* (Fig. 3D; Table 3) but not for *L. maculans* (Fig. 2D; Table 3).

### Effect of pH and Eh in the agar media on fungal growth

The addition of electrolyzed water products to malt-agar media extended the pH and Eh range on both sides of the native medium (Fig. 4A) and on both sides of the range from 5.5 to 6.5 observed in oilseed rape stems (Fig. 2, 3). The Eh-pH couples obtained did align on the theoretical correlation between Eh and pH, with a negative slope of −59 mV/pH unit. The native rice flour medium was pH 6.5, higher than the native malt medium at pH 4.5 to 5. It was more reduced, with Eh 400 mV compared to Eh 550-600 mV. The addition of Hydrogen peroxide, and Hydroquinone to rice flour media allowed variation in Eh (from 300 to 500 mV), but less variation in pH (6 to 7) (Fig. 4B).

**Fig. 4.**
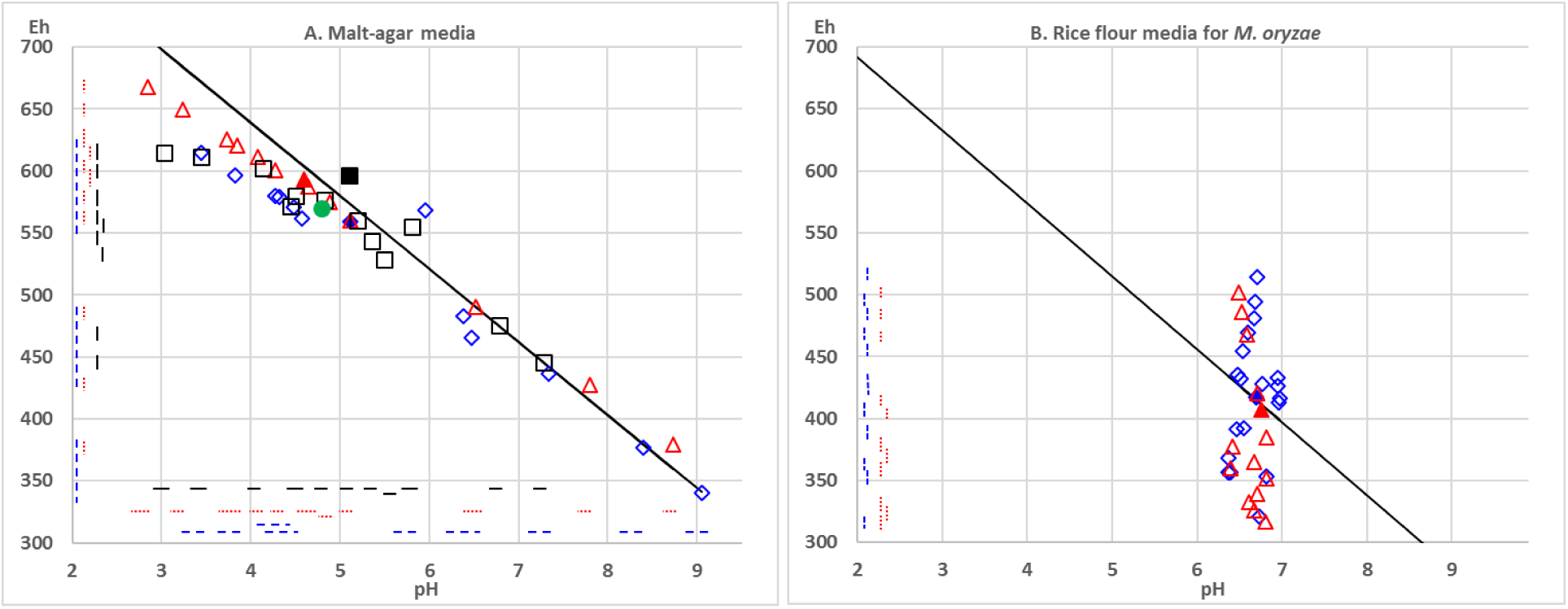
Range of pH and Eh in the native (filled symbols) and modified (empty symbols) media, with A. malt-agar media in four independent experiments Jan2015 (diamonds) Feb2015 (triangles) Jul2015 (squares) and Mar2016 (circle). B. rice flour-agar media in two independent experiments Jan2016 (diamonds) and Mar2016 (triangles). Line shows the theoretical relationship between pH and Eh, with a slope of −59 mV/pH unit according to the equation of Nernst. Dashed lines correspond to significantly different groups for pH (horizontal) and Eh (vertical) in the REGW-Q analysis after ANOVA (see Table 2).

For the six fungi tested, mean colony diameter varied on the range of 13 malt-agar media (Fig. 5; Table 2). Further, the colony appearance of all fungi, the pattern of sclerote production for *S. minor* and *S. sclerotiorum*, the production of yellow pigments for *L. biglobosa* varied in a manner congruent between replicated dishes in independent preparations (Supplementary information Fig. S3). In most cases, the fungal response to the modified media was quantitative with a decrease in colony diameter. However, in a few occasions the response was also qualitative, with the fungus either remaining on the transferred agar plug for some of the dishes, or starting to grow on some other dishes (Fig S3). The growth of *A. brassicicola, L. biglobosa* and *V. dahliae* was better at higher pH / lower Eh than at lower pH/higher Eh, with optimum pH around 7-8 and optimum Eh around 450 mV (Fig. 5). The growth of *L. maculans* was reduced on both sides of the range tested, with optimum pH around 6 and optimum Eh around 500 mV. The optimum were lower for *S. minor* with pH 5-6, and Eh below 400 mV than for *S. sclerotiorum* with pH around 6 and Eh around 450 mV.

**Fig. 5.**
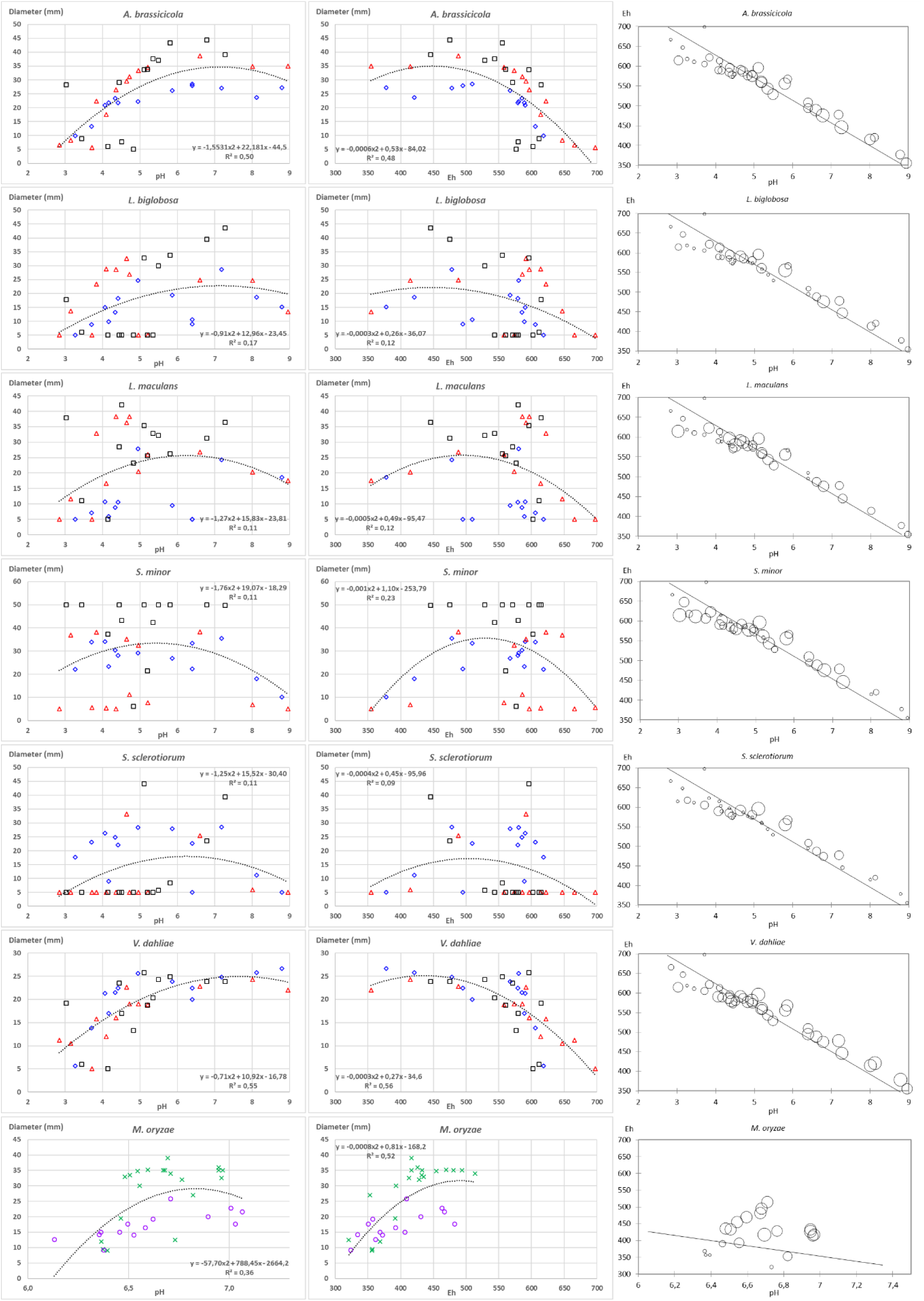
Mean colony growth of 6 fungi (each on one row of the figure) depending on pH (left panel), Eh (central panel), and the combination of pH and Eh (right panel) in the media prepared in three independent experiments Jan2015 (diamonds) Feb2015 (triangles) and Jul2015 (squares). Polynomial regression over all pH and Eh data is indicated. Each point is mean of 2 measures on 3 colonies (Jan2015, Feb2015) and 9 colonies (Jul2015). Corresponding Eh-pH ranges of the media are given in Fig.4. Note that pH X-axis scale is different for *M. oryzae*. Line in the right panel shows the theoretical relationship between pH and Eh, with a slope of −59mV/pH unit according to the equation of Nernst.

In the Sep15 experiment, the addition up to 500ppm of NaCl to the malt-agar media influenced Eh and pH (Fig. S1A). All media were more reduced (Eh from 480 to 495 mV) than the native one (Eh = 505mV). Compared to the native one (pH = 5.6), the pH was decreased (5.45 and 5.5) only for the addition of 100 and 500ppm of NaCl. The colony growth of the five fungi *S. minor, L. maculans, L. biglobosa, A. brassicicola* and *V. dahlia* was not altered by NaCl. The colony growth of *S. sclerotiorum* differed among media, though the only growth different from the native medium N0 was with 250 ppm of NaCl (Fig. S1B).

## Discussion

The growth of all fungi tested significantly altered either Eh, pH or both in agar media (Fig. 1). This effect was consistently observed among replicates within each experiment, and among independent experiments. Changes in pH were congruent with previous literature for fungi notorious to acidify (e.g. *Sclerotinia*) or alkalinize (e.g. *Magnaporthe, Botrytis*) the media (Landraud *et al*., 2013; Liu *et al*., 2017; Vylkova *et al*., 2017). Changes in Eh were congruent with previous literature, with various aerobes strongly reducing the culture media, due to the consumption of oxygen (Rabotnova & Schwartz, 1962). These authors mention that as opposed to organisms able to control their internal conditions, microorganisms maintain their pH-redox homeostasis by modifying the environment where they live. In our independent experiments with six fungal species, congruent results indicate that the protocol developed for measurements in soil (Husson *et al*., 2016) can also be used for measurement in agar media.

Eh and pH are inversely correlated, and this process can be explained because oxidation of two molecules of water (losing 4 electrons) produces O2 and 4H+, thus leading to acidification. From the Nernst equation, it can be calculated that the theoretical slope of this correlation is a 59 mV decrease in Eh for one unit increase in pH at 25°C (Husson *et al*., 2018). In our experiment, we observed that without biological activity, all the agar media prepared align on this theoretical line (Fig. 4). In contrast, the final measurements in agar media after fungal growth do not match this theoretical correlation line (Fig. 1). Rabotnova and Schwartz (1962) postulate that the observed reduction of the media is due to the consumption of dissolved oxygen. Further, the available nutrients also influence the outcome, as observed with *R. solani* acidifying the media in presence glucose (Boswell *et al*., 2003). Thus, further studies on media with different compositions are needed to investigate whether the final values depend on the consumption / release of different nutrient and compounds by the fungi, or if final values are stable, thus indicating regulation.

Interestingly, the joint characterization of pH and Eh provides discrimination among species not discriminated by the pH alone. Measuring Eh reveals that even the species not modifying pH can have an impact on the surrounding environment. For plant crop species, plant pathogens on crops are controlled to avoid deleterious effects on plant yield. However, visible damage to plants does not always explain the whole of the deleterious effects. Realizing that the plant pathogens might interfere with the plant oxido-reduction status (De Gara *et al*., 2003; Williams *et al*., 2011; Kabbage *et al*., 2015) and thus that maintaining redox homeostasis might come at a cost for an infected plant, opens another direction of investigation to better understand pathogen damage to plants.

In infected plant stems, pH was significantly altered, in opposite directions for *L. maculans* (Fig. 2) and *S. sclerotiorum* (Fig. 3). The observed alcalinisation or acidification correlates with canker length. The direction of this change is congruent with changes observed on agar media (Fig. 1). As the measurements in plants result both from fungal growth and from the plant reaction to infection, follow-up studies are needed to disentangle the respective roles of fungal mycelium, of fungal metabolism and of plant reaction (Casdeval & Pirofski, 2003). Nevertheless, our study already confirms that direct measurement of Eh and pH is possible on infected plants. In plants, the balance between oxidant and antioxidant pool sizes, plays signaling roles in the regulation of gene expression and protein function in a wide variety of plant physiological processes including stress acclimation (Shigeoka & Maruta, 2014; Choudhury *et al*., 2017), regulation of plant development processes, organogenesis, senescence and defense against pathogens (Nosek *et al*., 2015; Marschall & Tudzynski, 2016; Noctor *et al*., 2017). Being able to perform direct measurements on infected plants opens the prospect of characterizing more precisely the joint effects of combinations of stresses, thus studying their interrelation.

It is known that disease severity depends on plant organ age (Kranz, 1990; Benada, 2017; Torres *et al*., 2017) and some pathogens are only able to infect specific of plant organs. We provide values for pH and Eh in stem, because we used stem pathogens. However, follow up studies are needed to map the pH and Eh values in the different organs of the plant at a given time, or the change in these values in a given organ over time. Also, being able to perform direct measurements on a small amount of plant material will allow detailed studies to test whether the change is only local or systemic in the plant, for example infecting one organ and measuring at another location of the same organ or in neighboring organs.

Comparing fungal growth on media with a range of pH has already been performed (Daval *et al*., 2013, Lebreton *et al*., 2014). However, when chemical compounds are added to the growth medium, it is difficult to disentangle the effects via pH from directs effects (Bekker *et al*., 2009). We tried to minimize this problem by using electrolyzed water products, as NaCl was the only compound added. At low concentrations, NaCl promoted growth of *Botrytis cinerea* (Boumaaza *et al*., 2015), but at high concentrations reduced growth of both *B. cinerea* (Boumaaza *et al*., 2015) and bread spoilers *Penicillium roqueforti* and *Aspergillus niger* (Samapundo *et al*., 2010). At concentrations up to 500 ppm, NaCl did not alter fungal growth for five species (Fig. S1). The growth of *S. sclerotiorum* was affected, though our highest concentration was not high enough to reduce growth. With this study, we show that from two way of altering Eh and pH, adding electrolyzed water products produce combinations that align on the theoretical slope (Fig. 4A) whereas adding Hydrogen peroxide and Hydroquinone produce combinations that do not align on the theoretical slope (Fig. 4B). Choosing one or the other allows either exploring the joint variation of Eh and pH, or focusing on the variation of Eh within a narrower pH range.

On modified agar media, we observed that most fungi could grow over a broad pH range (Fig. 5) which is congruent with previous literature. Working mostly on bacteria, Rabotnova and Schwartz (1962) observed that microorganisms that have become adapted to special living conditions, for example pathogenic bacteria, grow only in narrow pH ranges. On the other hand, saprophytes that are diffused everywhere in nature can exist within wide pH ranges. We could see no such limitation within the species tested. Among the species tested, pH optimum was the highest for *V. dahliae* (Fig. 5) around pH 7 to 8. This fungus is able to grow hyphae in soil, which pH is less acidic that plant environment. For *A. brassicicola,* optimal pH between 5.5 to 7.5 was reported (Dinh *et al*., 2016). This is congruent with our results (Fig. 5). In *R. solani*, the pH of agar media modified the ability to solubilize metal phosphates, thus affecting growth of the colony (Jacobs *et al*., 2002). Beside the ability to grow, the environment conditions affect fungal virulence. This was documented in *B. cinerea*, for which having a functional thoredoxin system, a process enabling the maintenance of redox homeostasis, was required to achieve virulence (Viefhues *et al*. 2014). In *S. sclerotiorum*, oxalic acid is required for plant infection (Wang *et al*., 2016) and mediates the host redox environment (Williams *et al*., 2011; Kabbage *et al*., 2015). Conversely, ageing mycelium on artificial medium loses the ability to produce oxalic acid (Wang *et al*., 2016). We observed both that *S. sclerotiorum* strongly acidifies the medium (Fig. 1) and that fungal growth is reduced at pH below 6, being stopped at pH 3 (Fig. 5). Thus, the modification of the surrounding environment by the fungus might retroact on the fungus.

As compared to pH, information on the effect of redox potential on fungal growth is scarce. In our experiment, beside data for colony diameter (Fig. 5) it was striking to see, that colony aspect was repeatedly and thoroughly modified (Fig. S3). We noticed that pigment production, mycelium color and aspect were affected. We did not attempt to quantify these changes, and further studies are needed, but it is noteworthy that pH and Eh impacted a wide range of fungal physiological processes. This is congruent with previous reports on impacts on regulation and gene expression (Daval *et al*., 2013; Lehman *et al*. 2015; Jwa & Hwang, 2017). In our experiment, the amount and pattern of sclerote production was strongly affected in *S. minor*, and to a lesser extent in *S. sclerotiorum* (Fig. S3). This is congruent with previous reports that sclerotial differentiation may be induced by reactive oxygen species. It has been shown that biogenesis of sclerotia in *S. rolfsii* is correlated with elevated levels of lipid peroxidation, whilst certain antioxidants (hydroxyl radical scavengers) inhibit differentiation in *S. rolfsii* and in other plant pathogenic sclerotium-forming fungi such as *R. solani, S. minor* and *S. sclerotiorum* (Georgiou & Petropoulou, 2001).

As a conclusion, our series of experiments indicate that the procedure published for Eh and pH in soil (Husson *et al*., 2016; 2018) can be extended for measurement in agar media and in infected plants. Further, the joint characterization of both parameters opens the way to a more precise understanding of the impact of fungi on their environment, and conversely, of the environment on fungal growth. This could have applications at the interface between plant pathology, microbial ecology and plant physiology. Indeed, disease severity interacts with organ age (Kranz, 1990) or senescence (Häffner *et al*., 2015) or nutrient status (Veresoglou *et al*., 2013). Contrasting disease outcomes have been described when different fungi grow on co-infected plants (Lamichhane & Venturi, 2015; Abdullah *et al*., 2017). The response of microbial communities or of biocontrol agents to pH have been documented (Singh *et al*., 2014; Chapelle *et al*., 2016). Furthermore, the processes of stress perception when pH is altered are being unraveled in yeast (Riback *et al*., 2017). However, a comprehensive understanding of these interactions or combinations of stresses is lacking (Nejat & Mantri, 2017; Pandey *et al*., 2017). Publications at the molecular level are congruent with an implication of oxido-reduction processes (Shigeoka & Maruta, 2014; Nosek *et al*., 2015; Marschall & Tudzynski, 2016; Choudhury *et al*., 2017; Noctor *et al*., 2017). These lines of evidence points to pH and Eh as key signaling factors and hub of stress in the plant-pathogen interactions. The availability of methods for measurement opens the prospect to study combinations of stresses, either on agar media or in plants, and get an understanding of the involvement of pH and Eh modifications in these interactions.

## Acknowledgements

All authors declare the absence of conflicts of interest. This work benefited from the financial support of INRA – the French National Institute for Agronomical Research.

## Supporting information

**Fig. S1.**
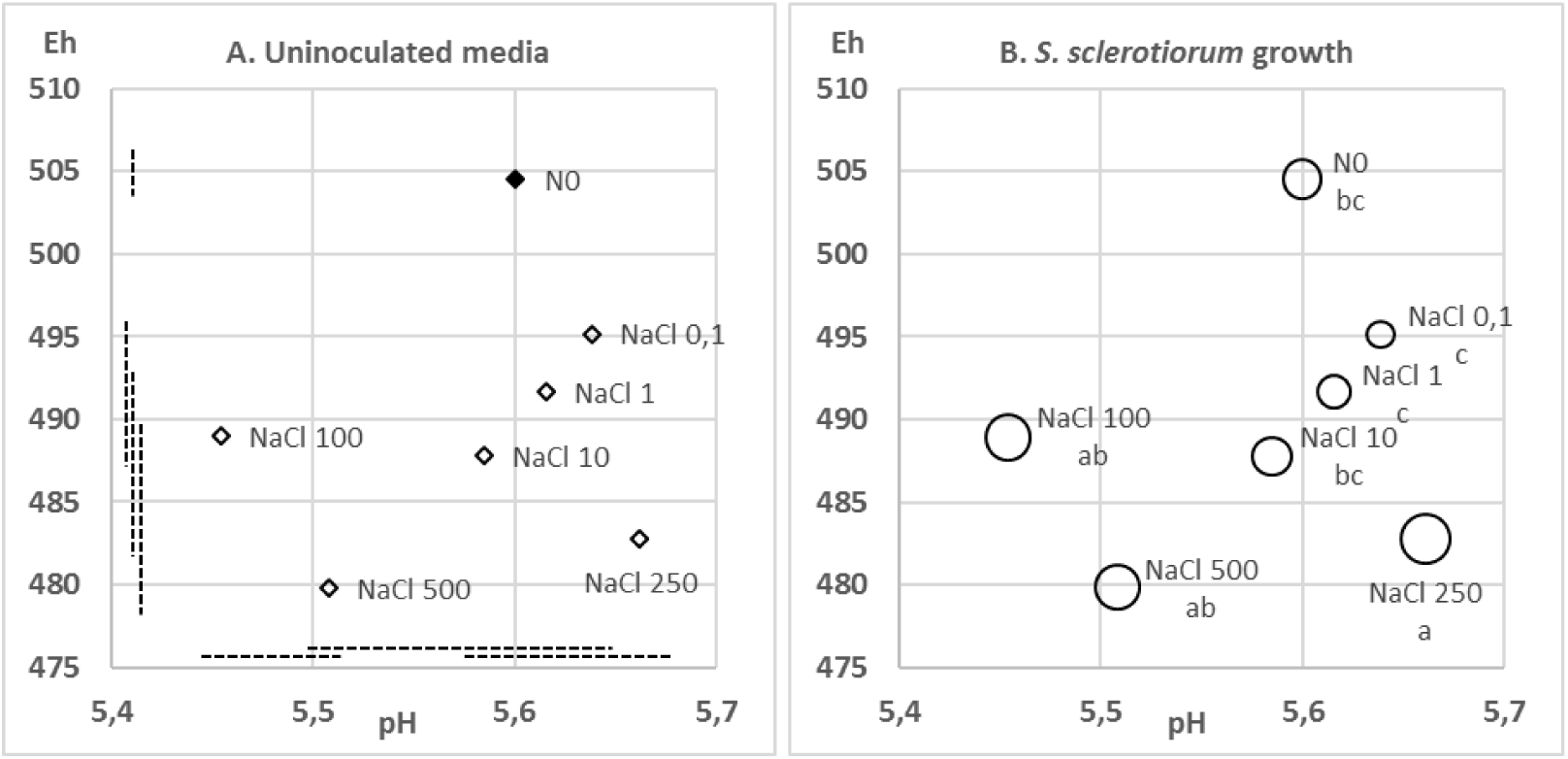
Effect of NaCl in the Sep15 experiment A. on the pH and Eh of the uninoculated malt-agar media; B. on the growth *S. sclerotiorum* colonies. Uninoculated medium (Panel A) was native (filled symbol) or with addition of NaCl up to 500 ppm (empty symbols). Dashed lines correspond to significantly different groups for pH (horizontal) and Eh (vertical) in the REGWQ analysis. In Panel B, size of circles correspond to colony diameter. Letters correspond to significantly different groups in the REGWQ analysis.

The experiment was performed on a range of 7 malt-agar media (Malt extract 20 g.l^-1^, Agar 20 g.l^-1^, Streptomycin 0.1 g.l^-1^) with increasing concentrations of NaCl (0, 0.1, 1, 10, 100, 250, 500ppm). Given the amount of NaCl added during water electrolysis and subsequent media preparation, final concentration was below 500ppm. Media were prepared 21/09/2015, inoculated 22/09/2015 and final assessment of colony growth was performed 25/09/2015 for *A. brassicicola, S. minor, S. sclerotiorum* and 08/10/2015 for *L. biglobosa, L. maculans* and *V. dahliae*.

**Fig. S2.**
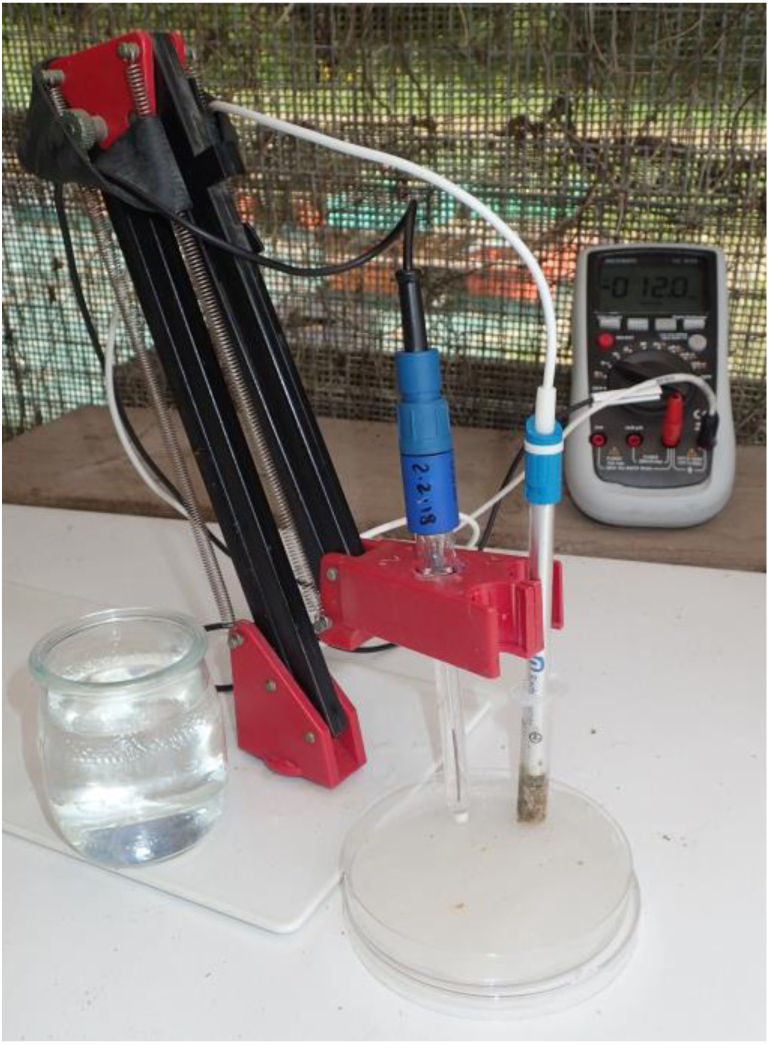
Measurement of Eh on plant stems. The plant material grinded with a mortar and pestle was placed in the barrel of a 5 ml syringe from which the nozzle had been cut. In the 9 cm Petri dish, the filter paper was moistened with 0.1M KCl solution. The Ag/AgCl reference electrode was placed standing, with tip touching the moist filter paper. The Pt plate electrode was placed in the plant material in the syringe, then the syringe barrel was placed standing with tip touching the moist filter paper.

**Fig. S3.**
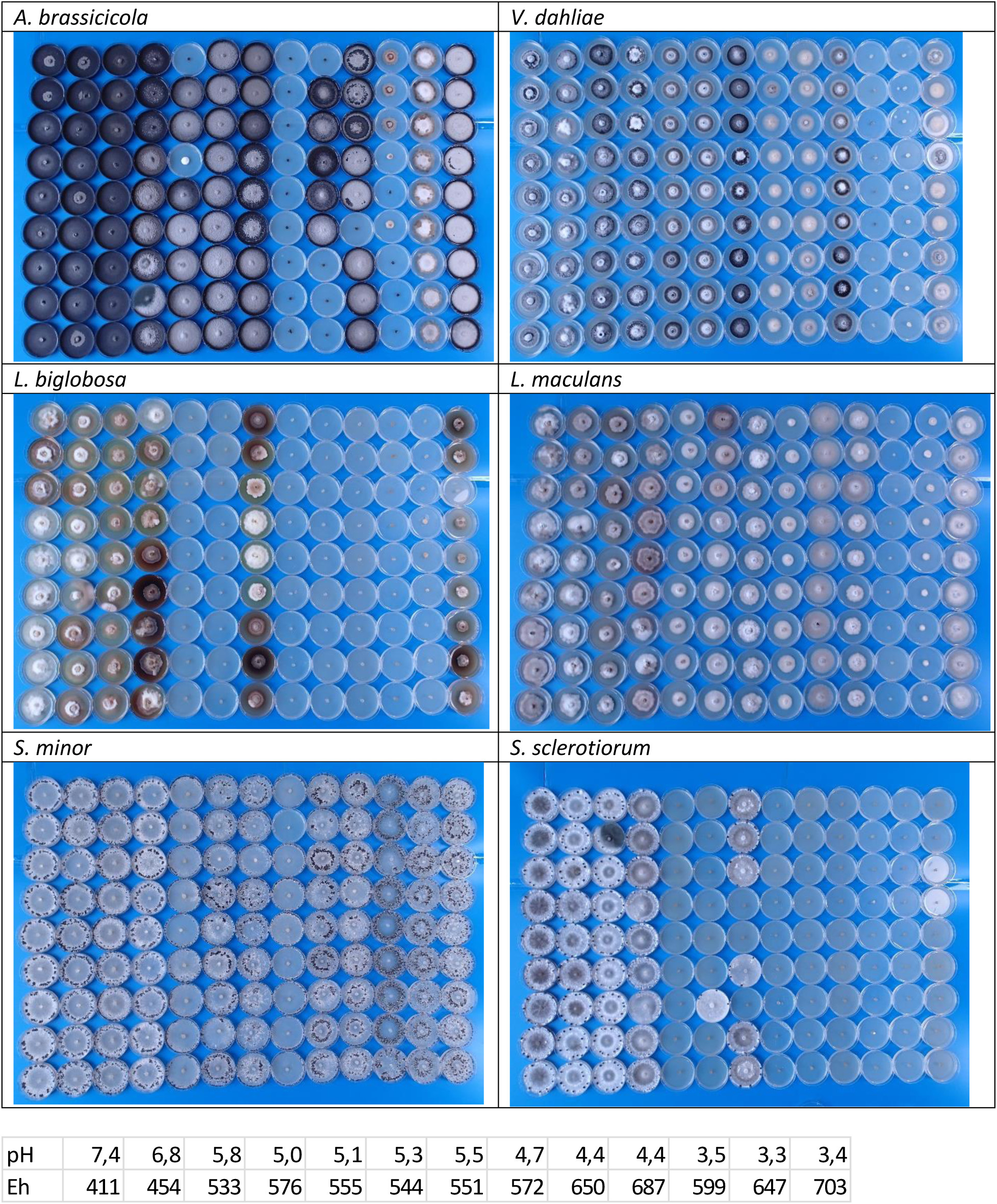
Colony aspect at the end of the Jul15 experiment on native and altered agar media for six fungi. Each panel correspond to a single strain transferred the same day and grown in the same room, with the 13 media in columns and replicates in lines. Within the experiment, lines 1 to 3; 4 to 6; 7 to 9 correspond to the three independent replicates (i.e. is preparation in independent bottles, and inoculation with agar plugs from a different source plate). Within upper, central and lower groups, dishes were prepared from the same bottles for all fungi. Mean pH and Eh values measured on aliquots corresponding to the 13 columns are given below. Note: colony diameter had been assessed earlier than the picture for fast growing fungi. Dishes are 5 cm in diameter.

## References

Abdullah A.S., Moffat C.S., Lopez-Ruiz F.J., Gibberd M.R., Hamblin J., Zerihun A. (2017) Host–multipathogen warfare: pathogen interactions in co-infected plants. Frontiers in Plant Science, 8, 1806.

Addinsoft (2016) XLSTAT: Data analysis and statistics with Microsoft Excel. Paris, France.

Bekker T.F., Kaiser C., Labuschagne N. (2009) The antifungal activity of potassium silicate and the role of pH against selected plant pathogenic fungi in vitro. South African Journal of Plant Soil, 26, 55–56.

Benada J. (2017) Measurement of redox potential and pH in plants and their function in the mechanism of plant resistance and in plant physiology. International Journal of Advanced Research in Electrical, Electronics and Instrumentation Engineering, 6, 1111–1116. doi:10.15662/IJAREEIE.2015.0501001

Boswell G.P., Jacobs H., Davidson F.A., Gadd G.M., Ritz K. (2003) Growth and function of fungal mycelia in heterogeneous environments. Bulletin of Mathematical Biology, 65, 447–477.

Boumaaza B., Benkhelifa M., Belkhoudja M. (2015) Effects of two salts compounds on mycelial growth, sporulation, and spore germination of six isolates of Botrytis cinerea in the western north of Algeria. International Journal of Microbiology, Article ID 572626, doi:10.1155/2015/572626.

Braunsdorf C., Mailänder-Sánchez D., Schaller M. (2016) Fungal sensing of host environment. Cellular Microbiology, 18, 1188–1200.

Casadevall A., Pirofski L.A. (2003) The damage–response framework of microbial pathogenesis. Nature Reviews. Microbiology, 1, 17–24.

Chapelle E., Mendes R., Bakker P.A.H.M., Raaijmakers J.M. (2016) Fungal invasion of the rhizosphere microbiome. The ISME Journal, 10, 265–268.

Choudhury F.K., Rivero R.M., Blumwald E., Mittler R. (2017) Reactive oxygen species, abiotic stress and stress combination. The Plant Journal, 90, 856–867.

Daval S., Lebreton L., Gracianne C., Guillerm-Erckelboudt A.Y., Boutin M., Marchi M., Gazengel K., Sarniguet A. (2013) Strain-specific variation in a soilborne phytopathogenic fungus for the expression of genes involved in pH signal transduction pathway, pathogenesis and saprophytic survival in response to environmental pH changes. Fungal Genetics and Biology, 61, 80–9.

De Gara L., de Pinto M.C., Tommasi F. (2003) The antioxidant systems vis-à-vis reactive oxygen species during plant–pathogen interaction. Plant Physiology and Biochemistry, 41, 863–870.

Dietz K.J., Mittler R., Noctor G., (2016) Recent progress in understanding the role of reactive oxygen species in plant cell signaling. Plant Physiology, 171, 1535–1539.

Dinh V.T., Somasekhara Y.M., Shivaprakash M.K. (2016) Cultural and physiological studies of Alternaria brassicicola (Schw.) Wiltshire of cabbage (Brassica oleracea var. capitata L.). Mysore Journal of Agricultural Sciences, 50, 529–534.

Foyer C.H., Noctor G. (2005). Redox homeostasis and antioxidant signaling: a metabolic interface between stress perception and physiological responses. The Plant Cell, 17, 1866–1875.

Foyer C.H., Noctor G. (2016) Stress-triggered redox signalling: what’s in prospect? Plant, Cell and Environment, 39, 951–964.

Georgiou C.D., Petropoulou K.P. (2001) Effect of the antioxidant ascorbic acid on sclerotial differentiation in Rhizoctonia solani. Plant Pathology, 50, 594–600.

Häffner E., Konietzki S., Diederichsen E. (2015) Keeping control: the role of senescence and development in plant pathogenesis and defence. Plants, 4, 449–488.

Howell D.C. (2009) Statistical methods for psychology. Cengage Learning. Wadsworth

Husson O. (2013) Redox potential (Eh) and pH as drivers of soil/plant/microorganism systems: a transdisciplinary overview pointing to integrative opportunities for agronomy. Plant and Soil, 362, 389–417.

Husson O., Brunet A., Babre D., Charpentier H., Durand M., Sarthou J.P. (2018) Conservation Agriculture systems alter the electrical characteristics (Eh, pH and EC) of four soil types in France. Soil and Tillage Research, 176, 57–68.

Husson O., Husson B., Brunet A., Babre D., Alary K., Sarthou J.P., Charpentier H., Durand M., Benada J., Henry M. (2016) Practical improvements in soil redox potential (Eh) measurement for characterisation of soil properties. Application for comparison of conventional and conservation agriculture cropping systems. Analytica Chimica Acta, 906, 98–109.

Jacobs H., Boswell G.P., Harper F.A., Ritz K., Davidson F.A., Gadd G.M. (2002) Solubilization of metal phosphates by Rhizoctonia solani. Mycological Research, 106, 1468–1479.

Jwa N.S., Hwang B.K. (2017) Convergent evolution of pathogen effectors toward reactive oxygen species signaling networks in plants. Frontiers in Plant Science, 8, 1687.

Kabbage M., Yarden O., Dickman M.B. (2015) Pathogenic attributes of Sclerotinia sclerotiorum: Switching from a biotrophic to necrotrophic lifestyle. Plant Science, 233, 53–60.

Kranz J. (1990) Fungal disease in multispecies plant communities. New Phytologist, 116, 383-405.

Lamichhane J.R., Venturi V. (2015). Synergisms between microbial pathogens in plant disease complexes: a growing trend. Frontiers in Plant Science, 6, 385.

Landraud P., Chuzeville S., Billon-Grande G., Poussereau N., Bruel C. (2013) Adaptation to pH and role of PacC in the rice blast fungus Magnaporthe oryzae. PLoS ONE 8(7): e69236.

Lebreton L., Daval S., Guillerm-Erckelboudt A.Y., Gracianne C., Gazengel K., Sarniguet A. (2014) Sensitivity to pH and ability to modify ambient pH of the take-all fungus Gaeumannomyces graminis var. tritici. Plant Pathology, 63, 117–128.

Lehmann S., Serrano M., L’Haridon F., Tjamos S.E., Metraux J.P. (2015) Reactive oxygen species and plant resistance to fungal pathogens. Phytochemistry, 112, 54–62.

Liu J., Zhang Y., Meng Q., Shi F., Ma L., Li Y. (2017) Physiological and biochemical responses in sunflower leaves infected by Sclerotinia sclerotiorum. Physiological and Molecular Plant Pathology, 100, 41– 48.

Lo Presti L., Lanver D., Schweizer G., Tanaka S., Liang L., Tollot M., Zuccaro A, Reissmann S., Kahmann R. (2015) Fungal effectors and plant susceptibility. Annual Review of Plant Biology, 66, 513–545.

Marschall R., Tudzynski P. (2016) Reactive oxygen species in development and infection processes. Seminars in Cell and Developmental Biology, 57, 138–146.

Nejat N., Mantri N. (2017) Plant immune system: crosstalk between responses to biotic and abiotic stresses the missing link in understanding plant defence. Current Issues in Molecular Biology, 23, 1–16.

Noctor G., Reichheld J.P., Foyer C.H. (2017) ROS-related redox regulation and signaling in plants. Seminars in Cell and Developmental Biology, 80, 3–12.

Nosek M, Kornaś A., Kuźniak E, Miszalski Z. (2015) Plastoquinone redox state modifies plant response to pathogen. Plant Physiology and Biochemistry, 96, 163–170.

Pandey P., Irulappan V., Bagavathiannan M.V., Senthil-Kumar M. (2017) Impact of combined abiotic and biotic stresses on plant growth and avenues for crop improvement by exploiting physio-morphological traits. Frontiers in Plant Science, 8, 537.

Prusky D., Yakoby N. (2003) Pathogenic fungi: leading or led by ambient pH? Molecular Plant Pathology, 4, 509–516.

R Core Team (2013) R; a language and environment for statistical computing. R Foundation for Statistical Computing, Vienna, Austria. URL http://www.R-project.org/.

Rabotnova I.L., Schwartz W. (1962). The importance of physical-chemical factors (pH and rH2) for the life activity of microorganisms. Microbiology. VEB Gustav Fisher Verlag, Jena.

Riback J.A., Katanski C.D., Kear-Scott J.L., Pilipenko E.V., Rojek A.E., Sosnick T.R., Drummond D.A. (2017) Stress-triggered phase separation is an adaptive, evolutionarily tuned response. Cell, 168, 1028–1040.

Rousk J., Baath E., Brookes P.C., Lauber C.L., Lozupone C., Caporazo J.G., Knight R., Fierer N. (2010) Soil bacterial and fungal communities across a pH gradient in an arable soil. The ISME Journal, 4, 1340–51.

Samapundo S., Deschuyffeleer N., Van Laere D., DeLeyn I., Devlieghere F. (2010) Effect of NaCl reduction and replacement on the growth of fungi important to the spoilage of bread. Food Microbiology, 27, 749–756.

Shigeoka S., Maruta T. (2014) Cellular redox regulation, signalling, and stress response in plants. Bioscience Biotechnology and Biochemistry, 78, 1457–70.

Singh A., Shahid M., Srivastava M., Pandey S., Sharma A., Kumar V. (2014) Optimal physical parameters for growth of Trichoderma species at varying pH, temperature and agitation. Virology and Mycology, 3: 127.

Smiley R.W. (1974) Take-all of wheat as influenced by organic amendments and nitrogen fertilizer. Phytopathology, 64, 822–825.

Stiles C.M., Murray T.D. (1996) Infection of field-grown winter wheat by Cephalosporium gramineum and the effect of soil pH. Phytopathology, 86, 177–183.

Tardi-Ovadia R., Linker R., Tsror L. (2017) Direct estimation of local pH change at infection sites of fungi in potato tubers, Phytopathology, 107, 132–137.

van der Does H.C., Rep M. (2017). Adaptation to the host environment by plant-pathogenic fungi. Annual Review of Phytopathology, 55, 427–450.

Veresoglou S.D., Barto E.K., Menexes G. Rillig M.C. (2013) Fertilization affects severity of disease caused by fungal plant pathogens. Plant Pathology, 62, 961–969.

Viefhues A., Heller J., Temme N., Tudzynski P. (2014) Redox systems in Botrytis cinerea: Impact on development and virulence. Molecular Plant Microbe Interactions, 27, 858–874.

Vylkova S. (2017) Environmental pH modulation by pathogenic fungi as a strategy to conquer the host. PLoS Pathogens, 13(2): e1006149.

Wang J.P., Xu Y.P., Zhang X.P., Li S.S., Cai X.Z. (2016) Sclerotinia sclerotiorum virulence is affected by mycelial age via reduction in oxalate biosynthesis. Journal of Integrative Agriculture, 15, 1034– 1045.

Williams B., Kabbage M., Kim H.J., Britt R., Dickman M.B. (2011) Tipping the balance: Sclerotinia sclerotiorum secreted oxalic acid suppresses host defenses by manipulating the host redox environment. PLoS Pathogens, 7, Doi=10.1371/journal.ppat.1002107.

Zeilinger S., Gupta V.K., Dahms T.E.S., Silva R.N.,Singh H.B., Upadhyay R.S., Vieira Gomes E., Tsui C.K.M., Nayak S.C. (2016) Friends or foes? Emerging insights from fungal interactions with plants FEMS Microbiology Reviews, 40, 182–207.

